# Laminar VASO fMRI in focal hand dystonia patients

**DOI:** 10.1101/2022.07.28.501870

**Authors:** Laurentius (Renzo) Huber, Panagiotis Kassavetis, Omer Faruk Gulban, Mark Hallett, Silvina G Horovitz

## Abstract

Focal Hand Dystonia (FHD) is a disabling movement disorder characterized by involuntary movements, cramps and spasms. It is associated with pathological neural microcircuits in the cortical somatosensory system. While invasive preclinical modalities allow researchers to probe specific neural microcircuits of cortical layers and columns, conventional functional magnetic resonance imaging (fMRI) cannot resolve such small neural computational units. In this study, we take advantage of recent developments in ultra-high-field MRI hardware and MR-sequences to capture altered digit representations and laminar processing in FHD patients. We aim to characterize the capability and challenges of layer-specific imaging and analysis tools in resolving laminar and columnar structures in clinical research setups.

We scanned N=4 affected and N=5 unaffected hemispheres at 7T and found consistent results of altered neural microcircuitry in FHD patients: a) In affected hemispheres of FHD patients, we found a breakdown of ordered finger representation in the primary somatosensory cortex, as suggested from previous low-resolution fMRI. b) In affected primary motor cortices of FHD patients, we furthermore found increased fMRI activity in superficial cortico-cortical neural input layers (II/III), compared to relatively weaker activity in the cortico-spinal output layers (Vb/VI).

Overall, we show that layer-fMRI acquisition and analysis tools are applicable to address clinically-driven neuroscience research questions about altered computational mechanisms at the spatial scales that were previously only accessible in animal models. We believe that this study paves the way for easier translation of preclinical work into clinical research in focal hand dystonia and beyond.

## 1. Introduction

Focal Hand Dystonia (FHD) is a disabling movement disorder characterized by involuntary movements, cramps and spasms. Common sub-forms are known as writer’s cramp and musician’s cramp (Karp 2017) with epidemiological prevalence up to 1 in 2500 people (Torres-Russotto and Perlmutter 2008). FHD is associated with pathological neural microcircuits in the cortical somatosensory system. FHD is expected to have closer (1-2mm) and more overlapping finger representation in the somatosensory system (Bara-Jimenez et al. 1998; Butterworth et al. 2003; Catalan et al. 2012; Elbert et al. 1998; Hallett 2011; Nelson et al. 2009). While invasive preclinical neuroimaging modalities in animal models allow researchers to probe specific neural microcircuits at the mesoscopic scale of cortical layers and columns (Goense et al. 2016), conventional non-invasive functional magnetic resonance imaging (fMRI) in humans cannot resolve such small neural computational units. Thus, until now, non-invasive research tools that are routinely applicable in patients, could not yet reach their full potential in translating pre-clinical findings of mesoscopic neural circuitry to patients. Recent developments in ultra-high field MRI hardware and functional contrast generation allows researchers for the first time to capture layer-specific fMRI responses at the spatial scale of sub-millimeter voxels. Specifically, mesoscale fMRI with blood volume sensitive VASO (Hua et al. 2013; Lu et al. 2003) methods can map functional activation changes in the laminar microcircuitry and columnar finger representations without unwanted large draining vein effects of conventional BOLD fMRI (Huber et al. 2017). For example, previous high-resolution VASO imaging could capture the fine scale finger movement representations across layers and columns in healthy volunteers (HV) (Huber et al. 2020; 2017). These proof-of-principle studies exploited highly optimized experimental environments that might not be fulfilled with patients (Fig. 1). Specifically, submillimeter fMRI in clinical populations as opposed to healthy volunteers might be challenged by the following constraints:

**Fig. 1:**
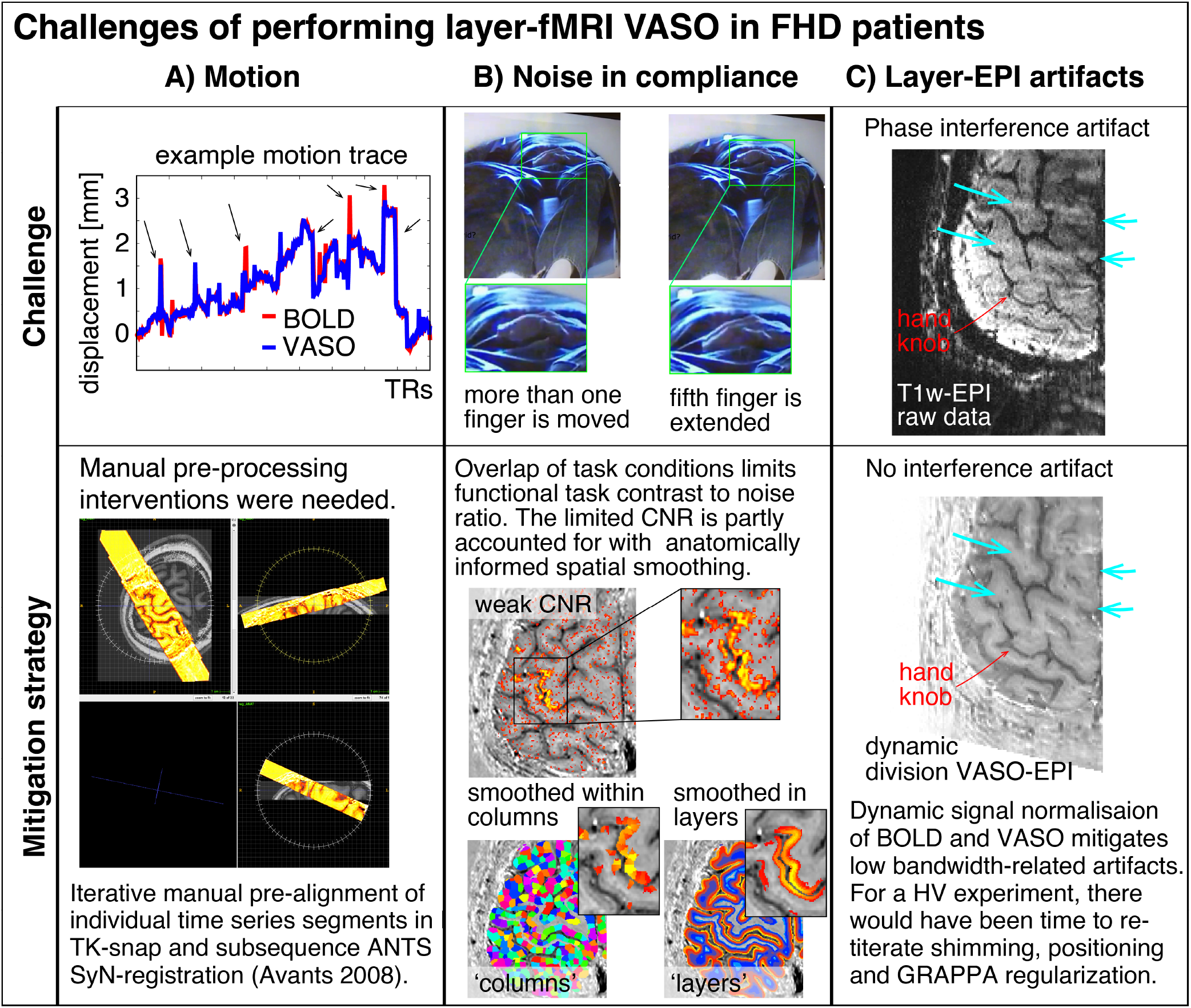
Selected examples of challenging aspects in the acquisition and processing of sub-millimeter VASO fMRI in FHD patients. In this study, head motion was mitigated with time consuming manual corrections. Limited finger specific functional contrast-to-noise-ratio (CNR) was mitigated with anatomically informed signal pooling within layers and columns, respectively. Time constant EPI phase interference artifacts were mitigated by means of dynamic division of odd and even time points with identical artifacts in SS-SI VASO.

- **Scan duration:** Previous studies could collect up to 18 hours of fMRI data per participant. This was achieved by inviting the participants for up to 10 two-hour scan sessions (Huber et al. 2020). Such large numbers of data collection sessions are not practical for FHD patients.
- **Head motion:** Access to patients is limited. Thus, the experimenter does not have the luxury to solely invite those individuals that have the ability to lie perfectly still for long periods of time (approx. 2h).
- **Noise in task compliance:** FHD patients can have a hard time following the task instructions. For example, when instructed to move the ring finger only, the middle finger and fifth finger are often moved too. This reduces the finger functional contrast-to-noise ratio.
- **EPI Artifact level:** Sub-millimeter fMRI is limited by low bandwidths and small coverages. In previous applications in healthy volunteers, this had been accounted for with iterative online fine tuning of acquisition parameters during the data acquisition phase (iterative B0-shimming, GRAPPA regularization, alignment, readout bandwidth). Within the limited scan time of patients, such fine tuning is not feasible.

Due to these challenges, the applicability of laminar and columnar functional imaging with fMRI in patient populations remains unclear. The aim of this study is to characterize and overcome these challenges. We used the altered digit representation in FHD patients as a testbed to explore the capability of applying layer-fMRI VASO to investigate mesoscopic neural representations in patients. Speficially, we sought to quantify pathological alterations of the mesoscopic finger representations in FHD patients across cortical depth as well as across topographical arrangements along the cortical sheet.

## 2. Methods

### 2.1. Patient procedure

Study participants were recruited under the NIH Institutional Review Board protocol 17-N-0126 (ClinicalTrials.gov identifier: NCT03223623). Four FHD patients (2M/2F;504.6y, disease duration 3 to 20y) receiving regular botulinum toxin injections were scanned at least 3 months after their last injection. Additionally, data from a prior study (Huber et al. 2020) were analyzed to control for task speed (N=2).

### 2.2. MRI scanner

The functional imaging sequence was implemented on a ‘classic’ MAGNETOM 7T scanner (Siemens Healthineers, Erlangen, Germany) using the vendor-provided IDEA environment (VB17A-UHF). For RF transmission and reception, a single-channel-transmit and 32-channel receive head coil (Nova Medical, Wilmington, MA, USA) was used. The scanner was equipped with a SC72 body gradient coil (maximum effective gradient strength used here: 49 mT/m; maximum slew rate used: 199 T/m/s).

### 2.3. MRI sequence

Slice-selective slab-inversion (SS-SI) VASO sequence (Huber 2014) was used in combination with a 3D-EPI readout, as previously implemented (Poser et al. 2010). The optimal sequence parameters were tested and optimized in previous studies (Huber et al. 2020). In short: No slab-oversampling, slab-excitation profile with a bandwidth-time-product of 25, T1-related blurring mitigation with variable flip angles, FLASH GRAPPA 3, vendor’s GRAPPA reconstruction algorithms (Siemens software identifier: IcePAT WIP 571), partial Fourier (6/8) reconstruction with POCS 8, in-plane resolution 0.75mm, slice-thickness: 0.89mm, TE=24ms, 24 slices, TR/TI=2.2/1.1s. A full list of protocol parameters is available on Github: https://github.com/layerfMRI/Sequence_Github/blob/master/FocalHandDystonia/FHD.pdf. The sequence binaries are freely available via the SIEMENS sequence ‘Appstore’ in Teamplay (https://teamplay.siemens.com/).

### 2.4. Scanning

The VASO imaging slab was positioned to be approximately perpendicularly oriented to the main surface of the central sulcus of one hemisphere. This was achieved by tilting it in two directions. Scanning was conducted using preset parameters for layer-fMRI VASO experiments, without individualized settings to optimize the time of the patients in the scanner. This scanning setup can be used to exemplify the usability and scalability of the experimental procedures beyond specialized MRI-development research groups. No specific custom hardware was necessary during the scanning, beyond the common 7T MRI configuration that is available at >100 centers worldwide. We scanned N=4 patients and N=2 healthy volunteers. Together, we collected data from N=5 healthy hemispheres and N=4 affected hemispheres.

### 2.5. Task

For the sake of consistency and comparability with previous studies, we used the same tapping tasks as previously used (Huber 2020). Briefly, before each run, subjects were instructed on what hand to tap with. Participants were instructed to tap one finger by extension-flexion at the metacarpophalangeal joint. The same tapping task was repeated for each finger. The tapping frequency was self-paced at a frequency of approximately every 1-2 seconds. During the scan, a video prompted which finger to move and when to rest. The tapping timing was locked to scanner triggers in units of 16 TRs (2.2s each) and contains visual cues when to tap which finger and for how long. The task was controlled via Psychopy 2 with publicly available scripts (https://github.com/layerfMRI/Phychopy_git/tree/master/Tapping_withTR_all_fingers). The same task was repeated for each hand. Each run lasted approximately 33min.

### 2.6. Processing

Motion corrected (ANTS)(Avants et al. 2008) data were sorted by contrast and corrected for BOLD contamination using LayNii v2.2.1(Huber et al. 2021) LN_BOCO. Layerification and columnification were done with the LN2_LAYERS program in LayNii. Block design activation z-scores and beta estimates were extracted with FSL-FEAT (Jenkinson et al. 2012). Cortical patch flattening was performed with the LN2_MULTILATERATE and LN2_PATCH_FLATTEN programs in LayNii (Gulban et al. 2021). Layer-extraction was manually constrained to the Brodmann area BA4a. This is the evolutionally older part of the primary motor cortex and located on the anterior side of the hand knob (also more lateral and closer to the skull as opposed to BA4p).

### 2.7. Re-analysis from previously published studies

We collected short video recordings of the FHD patients while they were performing the tapping task (example screenshots in Fig. 1B). This video material suggested a qualitative trend that FHD patients might have performed the tapping at a slower pace than what we were used to from previous studies with the same task. In order to quantify the effect of the tapping frequency on the interpretability of the results of this study, we obtained previously acquired layer-fMRI VASO data that were recorded with varying tapping frequencies. As described in (Huber et al. 2017), the resolution was 0.75mm x 0.75mm x 1.2mm, with VASO, at 7T in healthy participants. The tapping frequencies were 2Hz and 0.25Hz, which covers the range of tapping frequencies that we saw in FHD patients.

## 3. Results

Across all datasets, we find tapping induced activation both in M1 and in S1. Activation maps of all hemispheres in Fig. 2 shows that the experimental sub-millimeter VASO imaging setup can reliably capture neurally-induced functional cerebral blood volume (CBV) changes in patients across hemispheres and individuals. It can be seen how the activity is confined to gray matter (GM) without the sensitivity of large drain pial veins above the GM surface.

**Fig. 2:**
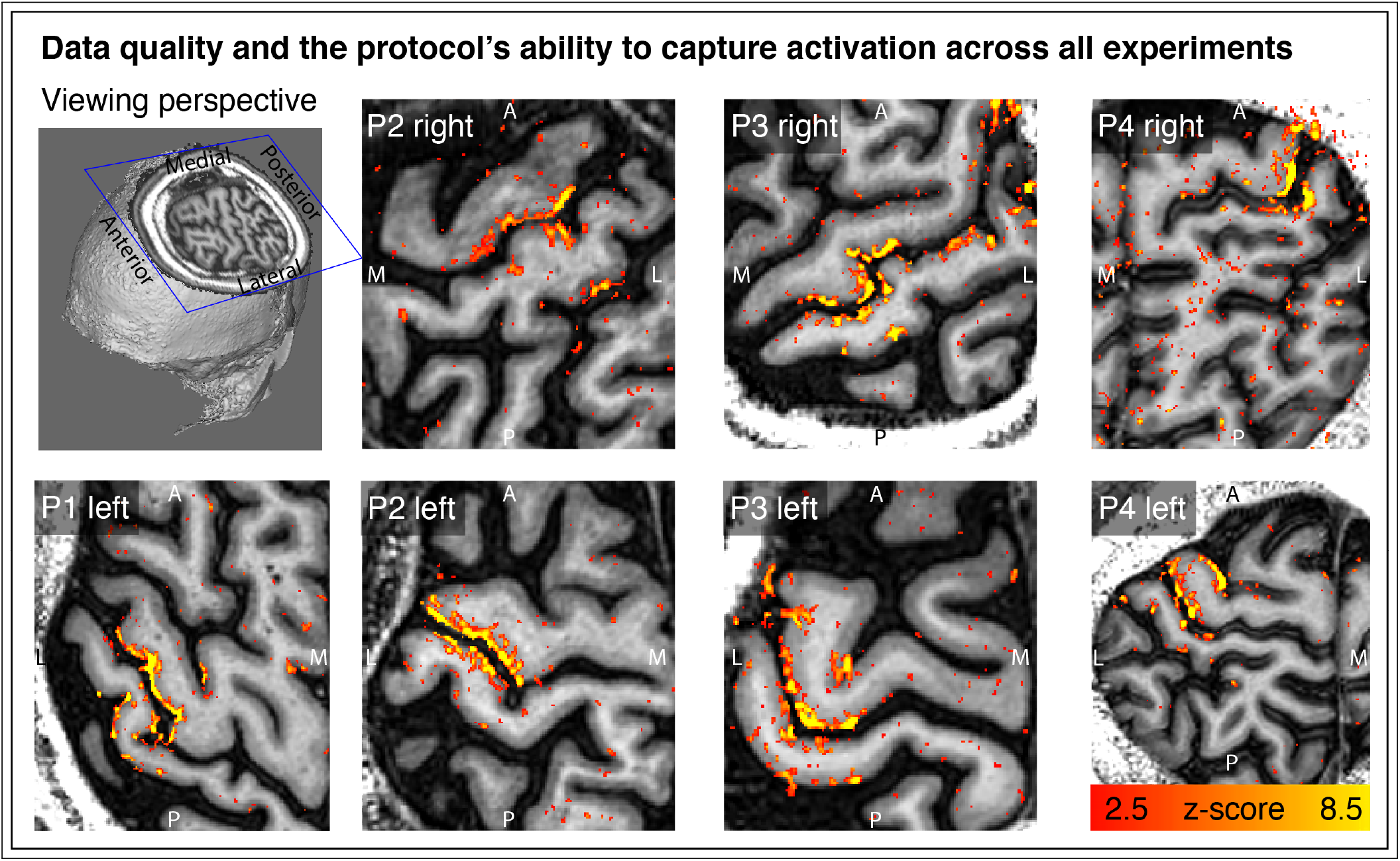
Tapping induced activation maps across patients. Within the 33min functional experiments, enough data are obtained to extract significant VASO signal changes across all patients and hemispheres. The figures represent the signal without spatial smoothing in spatially upsampled in-plane resolution of 0.4mm (nominal resolution 0.75mm).

In order to explore the spatial structure of finger representation in the primary somatosensory cortex on the posterior bank of the central sulcus, we imposed a local coordinate system within the GM cortical sheet. Exemplary data of one representative participant show exemplary how the finger representation follows the somatotopic alignment for the healthy hemisphere (Fig. 3). The spatial structure of the representations in the FHD-affected hemisphere are less clear. This is not for the lack of detection sensitivity. In fact, large patches of the primary somatosensory cortex are clearly engaged during finger tapping tasks. It’s just that the spatial arrangement of the finger-specific activation patches does not follow as clear arrangements as for the healthy hemisphere.

**Fig. 3:**
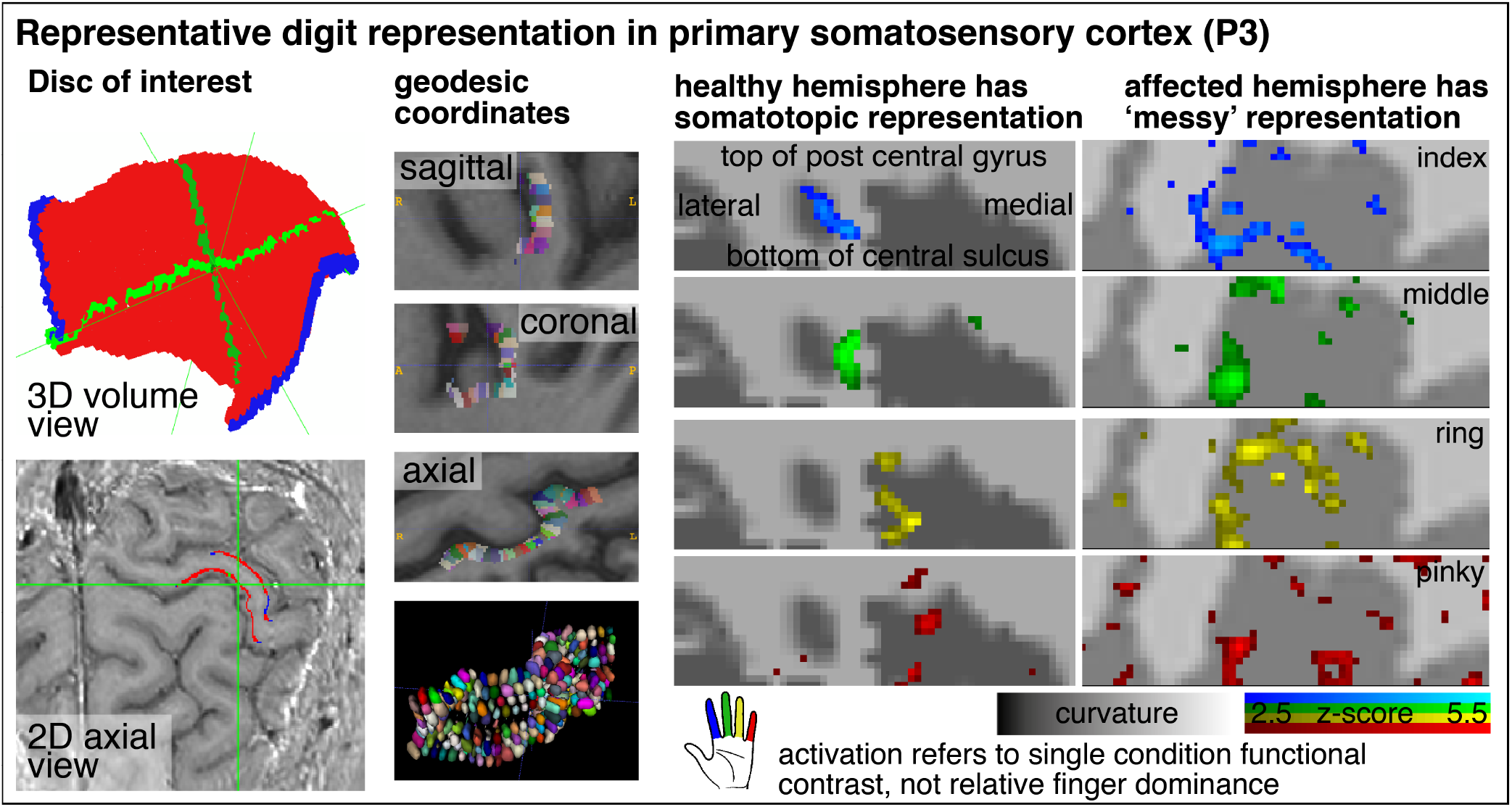
Cortical flattening of thin MRI slab. The first two columns exemplify how we flattened the cortical surface despite that the thin slab does not fulfill common topology requirements that are necessary in mesh-based analyses. Here, we imposed a local coordinate system in the distorted EPI data with `LN2_MULTILATERATE` (Gulban et al. 2021). Representative VASO-fMRI finger responses show somatotopic alignment in the healthy hemisphere and less so in the affected hemisphere (smoothed across layers only, no smoothing in the lateral direction).

Aside from topographical finger-representations in the sensory cortex, we also explored the laminar blood volume responses in the primary motor cortex. Fig. 4 depicts M1 layer-profiles of all patients. As expected, GE-BOLD signals show an increased bias towards the superficial layers compared to CBV-sensitive VASO. The affected hemispheres seem to be more dominated from superficial cortico-cortical input-layers compared to the healthy hemispheres, in all participants.

**Fig. 4:**
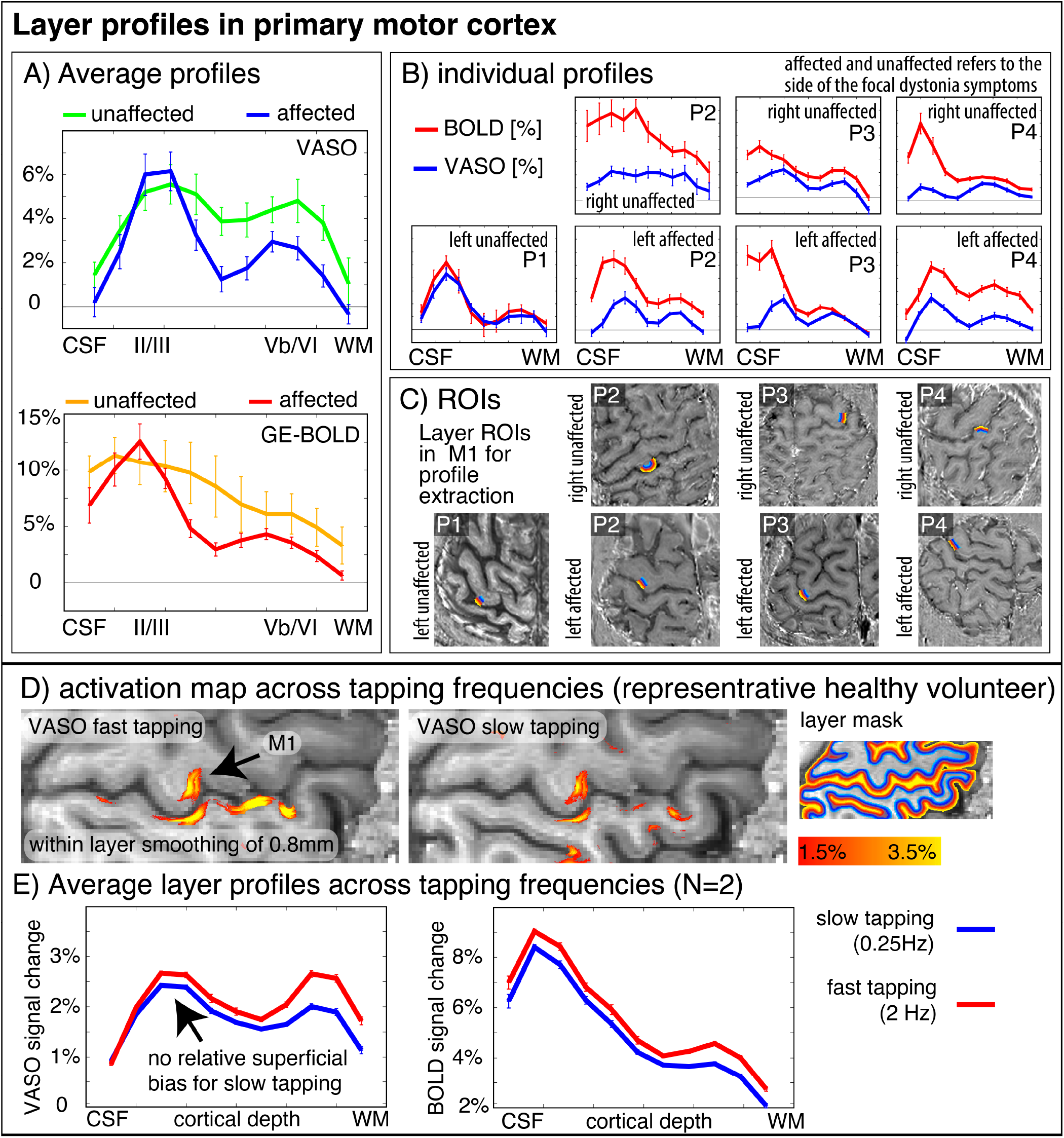
Layer-fMRI profiles in the primary motor cortex for BOLD and VASO in healthy and affected hemispheres. Panels A-B) highlight how all layer profiles consistently show a larger superficial bias in GE-BOLD compared to VASO, while still showing clear indication of a secondary “bump” in the deeper output layers. Compared to the deeper layers, the superficial layers in the affected hemispheres seem to be relatively stronger activated compared to their healthy counterparts. Signal pooling of layers was done from unsmoothed data. The corresponding ROIs are depicted in Panel C. Panels D-E) show layer-dependent CBV responses across tapping frequencies in healthy participants. It can be seen that slower tapping results in overall smaller fMRI signal changes. The relative reduction of fMRI signal changes is stronger in the deeper ‘output’ layers compared to the superficial ‘input’ layers. This suggests that the layer-specific signal modulation between healthy and affected hemispheres in Panel A cannot be explained by different tapping frequencies.

Video material of the FHD patients performing the task inside the scanner bore suggested a qualitative trend that FHD patients might have performed the tapping at a slower pace compared to typical tapping frequencies in healthy participants. To quantify the effect of tapping frequency on the interpretation of the layer profiles shown in Fig. 4A, we also show the tapping frequency dependence of the layer profile as collected in previous studies. These results are shown in Fig. 4E and suggest that a potentially different tapping frequency cannot explain the layer-dependent fMRI signal modulations that we find in healthy vs. affected hemispheres.

## 4. Discussion

In this study, we explored the capabilities and challenges of high-resolution layer-fMRI to inform research questions of affected cortical microcircuitry in patients. We scanned N=4 focal hand dystonia patients and found consistently altered somatotopic and laminar activation patterns in affected and unaffected hemispheres. The results presented here suggest that the high-resolution acquisition and analysis protocols developed here allow clinical neuroscientists to investigate pathological laminar pathologies in patients. Specifically, we found that cortical hemispheres that were affected by FHD showed a less structured somatotopic alignment of finger representation in the primary somatosensory homunculus of the postcentral gyrus. These results are consistent with the parallel ongoing imaging efforts of high-resolution functional mapping of finger representations at 7T (Pakenham et al. 2019). This study had already overcome some challenges and mapped the finger representation in FHD patients with 1.5mm GE-BOLD (Pakenham et al. 2019). Here, we pushed the imaging protocols further and used sub-millimeter veinbias-free VASO to capture laminar fMRI modulations. Aside from the altered somatotopic topographical digit representations in the primary sensory cortex, we also explored layer-dependent fMRI response in the primary motor cortex. We found that the superficial cortico-cortical input layers show increased activation during finger tapping tasks compared to unaffected hemispheres while the opposite is true for deeper layers. The increased activation may be due to decreased inhibition, and, in this way, these findings would be consistent with the notion that FHD is associated with a lack of neural surround-inhibition mechanisms (Hallett 2011). However, the relative dominance of fMRI signal changes in input layers could also be partly related to increased mental load (attention, motor planning, incorporating sensory feedback) and corresponding cortico-cortical input that is expected in the superficial layers (Persichetti et al. 2020; Trampel et al. 2012; Turner 2016). While the tapping frequency might have been slower for affected and unaffected hemispheres, we do not think that this can influence the interpretability of the layer-dependent activity modulation. In fact, slower tapping frequencies are expected to result in weaker superficial activation (Fig. 4E), which we do not see in the results of FHD affected hemispheres.

### 4.1. Relevance of this work

Looking beyond FHD, the tools developed and tested in this study provide a starting point for mapping layer-specific connections in patient populations in the general context of clinical neuroscience. Many influential theories of brain function of disorders posit hypotheses of neural deficits in distinct cortical layers and their role of feedforward and feedback processing. For example: mental disorders, such as autism and schizophrenia, and neurodegenerative diseases such as Parkinson’s or Huntington’s disease. Specifically:

- Layer-fMRI VASO has been discussed to be the key future technology to probe predictions of axonal loss and microcircuit dysfunction to provide insights for neurodegenerative motor disease (McColgan et al. 2020; Schreiber et al. 2021). The results shown there suggest that the imaging and analysis methodology is now ready to be applied for such proposed studies.
- Layer-fMRI has been discussed to be a key future technology to act as Occam’s razor for multiple competing hypotheses of hierarchical microcircuit disruptions in psychosis (Haarsma et al. 2020; Stephan et al. 2019). Namely in the context of predictive coding, one theory about psychosis discusses delusion symptoms in terms of deficits in learning. This is associated with respective predicted pathological neural computations in feedback dominated superficial and deeper layers. Another theory of psychosis discusses the same hallucination symptoms in terms of deficits in perceptual inference with respective predicted pathological neural computations in feedforward dominated middle layers. Until now, the methodological ability to constrain the models with empirical data has been limited. The usability of the layer-fMRI VASO protocols developed here allows future clinical neuroscientists to test and thus constrain these models.

These examples show that layer-fMRI applications in patients open the door to investigating computational mechanisms at the spatial scales that were previously only accessible in animal models. We believe that this study paves the way for easier translation of preclinical work into clinical research in focal hand dystonia and beyond.

### 4.2. In conclusion

While there are plentiful review articles about the value of layer-fMRI for clinical neuroscience research (Haarsma et al. 2020; McColgan et al. 2020; Schreiber et al. 2021; Stephan et al. 2019) experimental layer-fMRI data are scarce (Wan et al. 2021). Here, we present fMRI results of laminar and columnar CBV changes in FHD patients. We could not confirm previous findings of closer finger representations (Bara-Jimenez et al. 1998; Catalan et al. 2012; Elbert et al. 1998) because the finger representations in our maps did not show clear locations from which the distances could be calculated. We find that FHD affected hemispheres had a relatively stronger activity in motor input-layers (II/III) compared to the activity in motor output-layers (Vb/VI). This might be due to reduced surround inhibition in the superficial layers of the affected hemispheres. It could also be related to increased mental load (attention, motor planning, incorporating sensory feedback) that is related to neural input terminating in superficial layers. More data are needed to ultimately determine the stability of this finding. This study has built the methodological groundwork that future clinically-focused neuroscience application studies of layer-fMRI can be based on.

## Acknowledgements

We thank Sean Marrett for bringing together members from HMCS, NINDS and from SFIM, NIMH. Being introduced by him was the starting point of this cross-institute, cross-discipline collaboration project. We thank David Linden for discussions on how to address potential differences of tapping frequency which resulted in additional validation experiments shown in Fig. 4D-E.

## Help with scanning

These data were acquired with the kind support of the FMRIF core facility, specifically with the friendly help from Sean Marrett to test the acquisition protocols used here. We thank Kenny Chung for scan support. We thank Benedikt Poser for kindly providing the 3D-EPI sequence code used here.

## Ethics

We thank Avanti Iyer for help with the IRB protocol 17-N-0126.

